# The FORCE panel: An all-in-one SNP marker set for confirming investigative genetic genealogy leads and for general forensic applications

**DOI:** 10.1101/2021.11.30.470354

**Authors:** Andreas Tillmar, Kimberly Sturk-Andreaggi, Jennifer Daniels-Higginbotham, Jacqueline Tyler Thomas, Charla Marshall

## Abstract

The FORensic Capture Enrichment (FORCE) panel is an all-in-one SNP panel for forensic applications. This panel of 5,422 markers encompasses common, forensically relevant SNPs (identity, ancestry, phenotype, X- and Y-chromosomal SNPs), a novel set of 3,931 autosomal SNPs for extended kinship analysis, and no clinically relevant/disease markers. The FORCE panel was developed as a custom hybridization capture assay utilizing ∼20,000 baits to target the selected SNPs. Five non-probative, previously identified World War II (WWII) cases were used to assess the kinship panel. Each case included one bone sample and associated family reference DNA samples. Additionally, seven reference quality samples, two 200-year-old bone samples, and four control DNAs were processed for kit performance and concordance assessments. SNP recovery after capture resulted in a mean of ∼99% SNPs exceeding 10X coverage for reference and control samples, and 44.4% SNPs for bone samples. The WWII case results showed that the FORCE panel could predict 1^st^ to 5^th^ degree relationships with strong statistical support (likelihood ratios over 10,000 and posterior probabilities over 99.99%). To conclude, SNPs will be important for further advances in forensic DNA analysis. The FORCE panel shows promising results and demonstrates the utility of a 5,000 SNP panel for forensic applications.

## 1. Introduction

Forensic DNA databases were built around the use of non-coding DNA markers for human identification, primarily short tandem repeat (STR) loci from autosomal and Y-chromosomal targets [1]. By using a core set of DNA markers with almost no significance outside the medico-legal system, forensic DNA profiling was set to evolve independently from, and in parallel to, other fields of genetics sans overlap. This technological independence was rapidly breached, however, with the adoption of investigative genetic genealogy (IGG) and high-density single nucleotide polymorphism (SNP) genotyping for solving high-profile criminal cases (e.g., the Golden State Killer) and for identifications of unknown human remains [2]. Extant direct to consumer (DTC) SNP databases provided an exponentially growing, vast resource that could produce a genetic trail to anyone whose relatives participated in the consumer DNA testing market and opted into genealogical DNA databases such as GEDmatch [3,4]. The IGG method involves determining an extended DNA profile from a crime scene (or other forensic) sample using high-density SNP genotyping microarrays [5] or whole-genome sequencing (WGS) [6]. Then the SNP genotype dataset is searched in a public DNA database to identify matches with significant DNA sharing, indicating a familial relationship. Genealogists reconstruct family trees from the matches, identifying branches that could represent possible suspects that are provided to law enforcement. Once the “investigative lead” is identified, the next phase in the DNA profiling reverts back to using the forensic DNA core markers (i.e., STRs) to confirm the identification. DNA from the person of interest is obtained for testing, and the suspect’s STR profile is compared to the STR profile produced from the crime scene sample. The final DNA-based identification rests on the court-admissible STR results, which constitute the definitive forensic evidence from the case.

Confirmatory STR testing is feasible when the DNA from a forensic sample is of standard forensic quality. Yet certain sample types contain degraded DNA making them unsuitable for STR profiling, most notably rootless hair shafts and aged skeletal remains. Furthermore, confirmatory STR testing is limited when direct/reference samples are lacking and distant relationships must be inferred, as in historical remains identifications. In such cases, mitochondrial DNA (mtDNA) analysis is often used to recover a DNA profile, as mtDNA is higher in copy number than nuclear DNA (nDNA); thus the chances of recovering a mtDNA target are higher. Since mtDNA is a non-recombining lineage marker whose DNA sequence is shared amongst maternal relatives, the utility of mtDNA is limited forensically because its discrimination power is fairly weak. Therefore, mtDNA best serves degraded DNA cases that can be bolstered by other types of evidence, DNA or otherwise. Recent methods for mitogenome massively parallel sequencing (MPS) have been applied to degraded DNA samples and validated for forensic use, including mini-amplicon PCR [7,8], primer extension capture (PEC) [9], and hybridization capture [10]. Such methods are very sensitive and capable of providing authentic mtDNA data from the most challenging of forensic specimens. These MPS workflows have also been adapted for SNP profiling of degraded DNA specimens [11-15]. Large SNP multiplexes suitable for extended kinship analysis can be readily enriched using custom hybridization capture [14]. However, such large panels may not be suitable for routine forensic casework due to the markers that may be linked and/or associated with pathogenic variants.

To provide an enrichment approach for the generation of a large number of SNPs from all sample types, the FORensic Capture Enrichment (FORCE) panel was designed as an all-in-one SNP panel for forensic applications. This panel encompasses all forensically relevant SNP markers (identity, ancestry, phenotype, X- and Y-chromosomal SNPs) and presents a novel kinship SNP set for distant relationship inference. The relatively small size of the FORCE panel minimizes the number of primers/probes per reaction to reduce the enrichment cost. The FORCE panel can be adapted to various enrichment methods, including hybridization capture, PEC, and possibly multiplexed PCR, making it a viable option for both high quality and degraded DNA samples. The enriched SNP targets can be sequenced on a variety of downstream MPS platforms. By limiting the number of SNPs, fewer MPS reads are required to obtain per-SNP coverage requirements and thus sequencing efficiency is maximized. The FORCE panel SNPs were sub-selected from SNPs on the genotyping microarrays used by DTC DNA testing companies, enabling cross-compatibility with externally developed SNP data. This is important for forensic validation purposes to demonstrate concordance and genotyping accuracy. Medically relevant SNPs were excluded from the FORCE panel design in order to mitigate genetic privacy issues around FORCE panel DNA databanks. Aside from general forensic sample use, the FORCE panel is ideally suited to confirm investigative leads gained with genetic genealogy that cannot be confirmed with STR typing, such as those from degraded DNA.

## 2. Materials and Methods

### 2.1 FORCE panel design

A kinship SNP panel was designed to provide maximum kinship resolution with minimal SNPs for enhanced sequencing efficiency. Kinship SNPs were selected based on the following criteria (Table 1): 1) inclusion in three commonly used Illumina SNP genotyping chips (Table 2; San Diego, CA), 2) minor allele frequency (MAF) between 0.2 and 0.8 in the major 1000 Genomes continental populations (African (AFR), Admixed American (AMR), East Asian (EAS), European (EUR) and South Asian (SAS)) [16], 3) at least 0.5 cM genetic distance between selected SNPs, 4) linkage disequilibrium (LD) metric r^2^ smaller than 0.1, and 5) frequency difference of less than 0.35 between the 1000 Genomes five continental populations. Using the same parameters as the autosomal kinship SNPs, X-SNPs were selected for cases requiring X-chromosomal information. Pre-existing SNP marker sets were added to the FORCE panel to offer cross-compatibility with commercial SNP assays (Table 2). Identity-informative SNPs (iiSNPs) were compiled from the ForenSeq DNA Signature Prep Kit (Primer Mix A; San Diego, CA), the Precision ID Identity Panel (Thermo Fisher Scientific, Waltham, MA), and a published QIAseq panel (QIAGEN) [17]. The ancestry- and phenotype-informative SNPs (aiSNPs and piSNPs, respectively) were compiled from the ForenSeq kit (Primer Mix B), the Precision ID Ancestry Panel (Thermo Fisher Scientific), and markers identified by the VISAGE consortium [18]. To increase the Y haplogroup resolution beyond the 34 Y-SNPs in the Precision ID Identify Panel, 884 Y-SNPs targeted by the amplicons of the Ralf *et al* [19] AmpliSeq panel (Thermo Fisher Scientific) were included in the assay. All of the selected FORCE panel SNPs were compared with American College of Medical Genetics (ACMG) secondary findings (SF) 2.0 genes to test for clinical relevance [20]. SNPs listed the ACMG SF 2.0 were omitted from the panel. Triallelic SNPs were omitted from the panel and autosomal microhaplotypes were not included to simplify the data analysis.

**Table 1.** Kinship SNP selection criteria and resulting viable SNPs.

| **Kinship Panel Criterion** | **Number of Remaining SNPs** |
| --- | --- |
| 1. Included on all major SNP array chips (Table 2) | 116,544 |
| 2. MAF 0.2-0.8 in 1000 Genomes populations | 34,851 |
| 3. Minimum 0.5 cM distance | 5,426 |
| 4. LD metric $r^2 < 0.1$ | 5,376 |
| 5. Maximum 35% frequency difference between 1000 Genomes populations | 3,937 |

**Table 2.** Commercial and/or published SNP panels utilized for the FORCE panel selection. iiSNP = identity-informative SNP; aiSNP = ancestry-informative SNP; piSNP = phenotype-informative SNP; X-SNP = X-chromosomal SNP; Y-SNP = Y-chromosomal SNP.

| Marker Type | SNP Panels (# SNPs) |
| --- | --- |
| Kinship SNP /<br>X-SNP | Infinium Global Screening Array (654,027), Infinium Omni Express (710,000), Infinium CytoSNP-850K (850,000) |
| iiSNP | ForenSeq DNA Signature Prep Kit: Primer Mix A (94) |
|  | Precision ID Identity Panel (90) |
|  | QIAseq Investigator 140 SNP panel (140) |
| aiSNP | ForenSeq DNA Signature Prep Kit: Primer Mix B (56) |
|  | Precision ID Ancestry Panel (165) |
|  | VISAGE panel (115) |
| piSNP | ForenSeq DNA Signature Prep Kit: Primer Mix B (24) |
|  | VISAGE panel (41) |
| Y-SNP | Precision ID Identity Panel (34) |
|  | AmpliSeq (884) |

A custom myBaits-20,000 kit (Arbor Biosciences, Ann Arbor, MI) was designed for hybridization capture of the selected SNPs. The design included four baits per autosomal SNP (right flank, left flank and each allele) and two baits per X/Y SNP (each allele). A BLAST [21] analysis was then performed to filter out non-specific baits. A total of 228 baits failed to meet the BLAST analysis requirements. Most of these failed baits (139/228) came from the AmpliSeq Y-SNP marker set [19], including 61 Y-SNPs for which both baits failed. The remaining failed baits targeted autosomal markers (74 SNPs with one failed bait, six SNPs with two failed baits, and one SNP with three failed baits). The failed baits were removed from the kit, thus eliminating 61 of the 884 Y-SNPs. Overall, there were 97 SNP markers with at least one failed bait and thus may show reduced coverage in the SNP capture results. The alpha version of the myBaits FORCE capture assay tested in this study included 5,446 SNPs (Table S1), targeted by 19,526 unique baits. There were 24 SNPs included in the alpha panel but ultimately excluded from analysis due to the outlined criteria. The excluded SNPs were six kinship SNPs with clinical relevance, 15 triallelic aiSNPs (one of which also had clinical relevance), one Y-SNP from the Precision ID Identity Panel with no Y haplogroup association, and two uninformative autosomal SNPs. A summary of the marker types comprising the 5,422 FORCE panel SNP targets analyzed in this study is presented in Figure 1 and detailed marker information is provided in Table S1.

**Figure 1.**
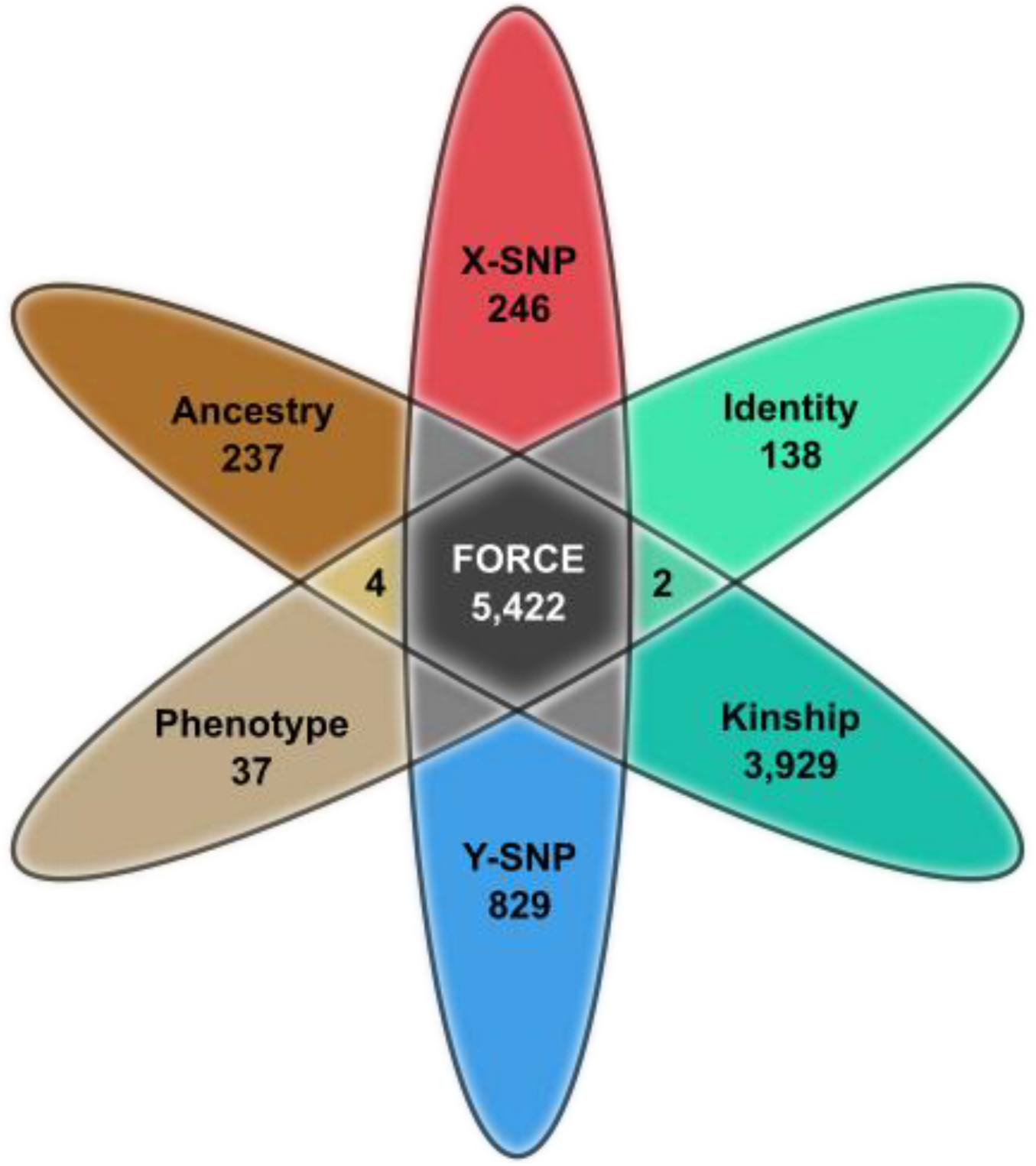
FORCE panel SNP marker summary (n=5,422).

### 2.2 Simulated Panel Performance

A simulation approach was used to study expected likelihood ratio (LR) distributions for various kinship case scenarios based on the kinship SNPs included in the FORCE panel. Five different kinship case scenarios were selected, representing 2nd to 6th degrees of pairwise relatedness (i.e. half siblings, first cousins, first cousins once removed, second cousins, and second cousins once removed). The simulations were performed with the complete kinship SNP set (the initial 3,937 SNPs including the six that were ultimately excluded). Simulations were also performed with 75% and 25% of the kinship SNPs in order to mimic situations for which only partial SNP profiles are available.

The simulations were performed using the software Merlin [22]. Merlin was used both to create the pedigree-based DNA data and to calculate likelihoods for the tested hypotheses. European allele frequencies from [16] were used and genetic linkage was accounted for using genetic position information from Rutgers map [23]. For each simulated kinship case scenario, the likelihood ratio was calculated as LR=Pr(DNA|H1)/Pr(DNA|H2), where H1 was the hypothesis representing the related case scenario, and H2 was the hypothesis representing the alternative, in which the tested individuals are unrelated. For each case scenario and true hypothesis, 10,000 simulation events were performed. A front-end script written in R [24] was used to format and manage input files, and R was also used to handle the outputs from Merlin and for plotting the distribution curves (R package *ggplot2*).

### 2.3 SNP Capture Assay Laboratory Testing

#### 2.3.1 Sample Selection

The samples utilized in this study originated from non-probative Defense Personnel Accounting Agency (DPAA) cases that were previously processed by the Armed Forces Medical Examiner System’s Armed Forces DNA Identification Laboratory (AFMES-AFDIL). These cases involve skeletal samples from five previously identified World War II (WWII) male service members and their associated family reference specimens. One or more DNA extracts from a single skeletal element per service member was included in this study, along with two or three family reference samples for each service member (Table 3). The five cases were chosen to range in sample quality and degree of relationship. In addition to the case samples included for testing the effectiveness of the kinship SNPs, further non-probative samples were evaluated for concordance and/or SNP performance (Table 4). These included seven reference samples, a 19^th^ century bone sample from a previously identified individual (sample 25, JB55 [12]), an additional 19^th^ century bone sample associated with JB55 (sample 26), and four control DNAs. All living sample donors provided informed consent for their samples to be used in research and quality improvement activities. The use of these reference and bone samples was approved by the Defense Health Agency Office of Research Protections (Protocol # DHQ-20-2073).

**Table 3.**
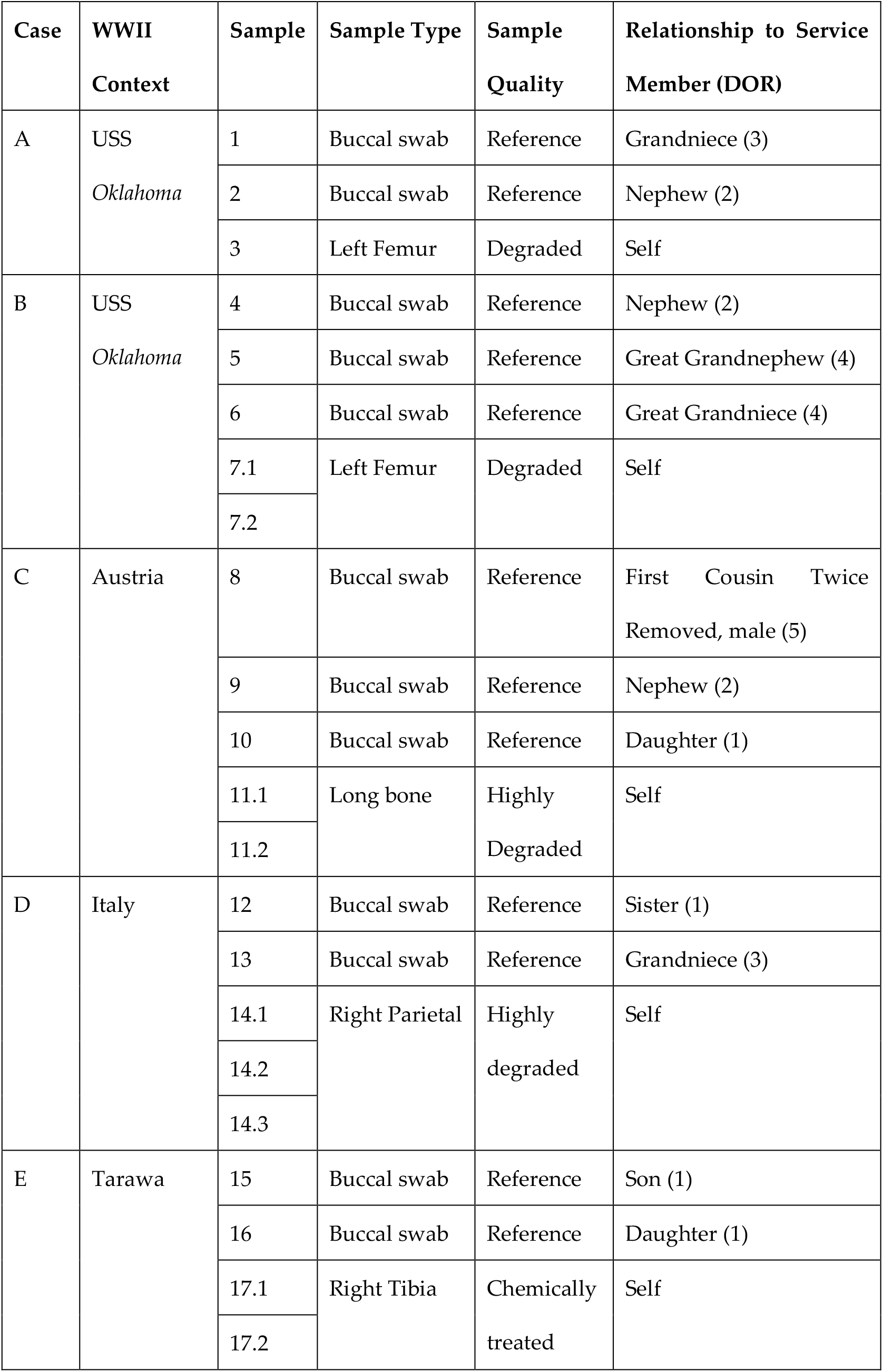
Non-probative case descriptions. Each of the five cases includes one or more DNA extracts from a bone sample of a previously identified World War II (WWII) service member, along with two or three family reference specimens. DOR=degree of relatedness.

**Table 4.** Additional non-probative samples tested.

| Sample | Sample Type | Sample Quality | Sex |
| --- | --- | --- | --- |
| 18 | Buccal swab | Reference | Female |
| 19 | Buccal swab | Reference | Female |
| 20 | Buccal swab | Reference | Male |
| 21 | Buccal swab | Reference | Female |
| 22 | Buccal swab | Reference | Female |
| 23 | Buccal swab | Reference | Male |
| 24 | Buccal swab | Reference | Male |
| 25 (JB55) | Femoral bone | Degraded (~200 years) | Male |
| 26 | Unknown bone | Degraded (~200 years) | Unknown |
| 2800M | Whole blood | Control DNA | Male |
| K562 | Cell line | Control DNA | Female |
| NA12877 | Cell line | Control DNA | Male |
| NA12878 | Cell line | Control DNA | Female |

#### 2.3.2 DNA Extraction and Quantitation

Existing DNA extracts and associated reagent blanks (RBs) from the five skeletal samples as well as JB55 [12] were obtained. During previous processing of these samples, DNA was extracted from 0.2-1.0 g bone powder. Samples were first digested overnight at 56°C in a buffer containing 0.5 M EDTA, 1% lauroylsarcosine and 200 µl of 20 mg/ml proteinase K. DNA extraction was achieved using either an organic protocol or a double QIAquick PCR Purification Kit method (QIAGEN, Hilden, Germany), as described in [25]. Purification was performed on all organic extracts using the QIAGEN MinElute PCR Purification Kit, followed by elution in 50 to 100 µl of Tris-EDTA [10 mM Tris, (pH 7.5) 0.1 mM EDTA]. An unknown skeletal sample from the same historical context as JB55, along with an associated RB, underwent a modified Dabney DNA extraction. This Dabney protocol followed the method using large volume silica columns [26]; however overnight digestion was performed at 56°C.

DNA was isolated from the buccal swabs using the QIAGEN EZ1 DNA Investigator Kit automated on an EZ1 Advanced XL instrument following the manufacturer’s Trace or Tip Dance Protocol. The final elution volume for all samples ranged from 50 to 100 µl of Tris-EDTA. DNA was quantified with the dsDNA High Sensitivity (HS) Assay Kit on the Qubit 2.0 or Qubit 3.0 Fluorometer (Thermo Fisher Scientific, Waltham, MA) to determine input into library preparation.

#### 2.3.3 Library Preparation

Samples were divided into three sets for laboratory processing and sequencing, and each set included library negative and positive controls to evaluate library and capture success. The first sample set (bone) included ten DNA extracts from the WWII bone samples and one of the historical bone samples (26) plus two associated RBs. The bone set positive control was 1 ng of fragmented K562 control DNA as previously described in [10]. JB55 was processed separately from the other bone samples in its own set, which included an associated RB, library negative control, and K562 positive control as used in the bone sample set. Lastly, the reference sample set included the 12 family references, seven additional reference-type samples, and three control DNAs (2800M, NA12877 and NA12878). Associated RBs and a library negative control were included in the reference sample set.

The bone libraries were prepared in a dedicated low copy DNA laboratory using the KAPA Hyper Prep Kit (Roche Sequencing, Wilmington, MA). Library preparation followed the manufacturer’s recommendations using 15 µM 8-bp dual-indexed adapters. A target DNA extract volume of 50 µL was used for library preparation, with a maximum DNA input of 1 µg. Some of the previously existing DNA extracts had limited volume available; therefore Tris-EDTA was added to create a total volume of 50 µL. Reference sample libraries were prepared with the KAPA HyperPlus Kit (Roche Sequencing), as fragmented DNA is required for successful library preparation. Library preparation was performed according to the manufacturer’s protocol with a 20-minute fragmentation and ligation of dual-indexed 8-bp adapters (Integrated DNA Technologies, Coralville, IA). The library PCR reactions for the bone samples utilized KAPA HiFi HotStart Uracil+ ReadyMix (Roche Sequencing) to accommodate for uracils present in the native DNA molecule from cytosine deamination. The reference/control sample libraries were amplified with the KAPA HiFi HotStart ReadyMix (Roche Sequencing). Library amplification was completed following the manufacturer’s recommendations, using a total of 12 cycles of PCR for each of the sample sets (bone and reference/control). Samples were purified using a 1.0X AMPure XP (Beckman Coulter, Indianapolis, ID) bead-based cleanup and were then eluted in 20 µl of Tris-EDTA. The quality of the libraries was checked on the 2100 Bioanalyzer instrument (Agilent Technologies, Santa Clara, CA) using the Agilent DNA 7500 Kit. For the bone sample and control DNA libraries, an aliquot of the purified library product was reserved for WGS (described below).

#### 2.3.4 Hybridization Capture

Hybridization capture with the alpha version of the FORCE capture panel was completed using the myBaits 1-20K Custom DNA Hybridization Capture kit (Arbor Biosciences, Ann Arbor, MI). The procedure followed the manufacturer’s recommended v5 protocol using a single capture and 5 µL of custom SNP baits. Samples were allowed to incubate overnight for approximately 24 hours at 62°C with a heated lid (72°C) using a Veriti thermal cycler (Thermo Fisher Scientific, Waltham, MA). PCR amplification was achieved using duplicate PCR reactions to make use of the entire capture product. The postcapture PCR reactions were completed with 19 PCR cycles using the KAPA HiFi HotStart ReadyMix following the manufacturer’s recommendations. The two amplified capture products were combined for each sample and then purified with the MinElute PCR Purification Kit, eluting in 25 µL of Tris-EDTA.

#### 2.3.5 Normalization, Pooling, and High Throughput Sequencing

Purified capture product was quantified using the 2100 Bioanalyzer instrument with the Agilent DNA 7500 Kit to determine DNA molarity. As in library preparation and hybridization capture, the three sample sets (bone, JB55 and reference) were pooled separately to minimize the impact of crosstalk between samples of disparate quality during sequencing [10]. The bone set positive control (K562) was omitted from the bone sample sequencing pool and instead added to the reference sample pool for sequencing. The JB55 sample pool included JB55 and its associated RB, as well as an additional library from a different JB55 DNA extract. (Since both JB55 libraries produced similar results; only the data from the DNA extract that also underwent WGS are presented below.) The JB55 library controls were not sequenced; but they were assessed with the Bioanalyzer for contamination and quality control purposes.

Pooling was achieved by individually normalizing captured libraries to the same concentration as the least concentrated sample (nM) in the set, then adding equal volume of all samples to the pool along with maximum RB and negative control volumes. The captured library pools were quantified using the Agilent DNA 7500 Kit on the 2100 Bioanalyzer. The bone and reference captured library pools were diluted to 1 pM with a 5% spike-in of PhiX Control V3 (Illumina) for sequencing on a NextSeq 550 (Illumina) using NextSeq 500/550 High Output Kits, v2.5 (150 cycles). Paired-end sequencing (75×75 cycles) was performed for the bone sample pool (14 captured libraries), while single-end data (150 cycles) were generated for the reference and control DNA samples (26 captured libraries). The JB55 pool was diluted to 12 pM including 5% PhiX control V3, and paired-end sequencing (75×75 cycles) was completed on a Verogen MiSeq FGx in RUO mode using a MiSeq Reagent Kit, v3 (150 cycles).

To gauge capture efficiency for the non-probative bone sample and control DNA libraries, baseline WGS data were produced on an Illumina NextSeq 550. WGS was not performed for any of the reference samples. Sequencing was completed with NextSeq 500/550 High Output Kits, v2.5 (150 cycles) using the same approach as the captured libraries (75×75 cycles for the bone sample pool and 1×150 cycles for the control DNA pool). For direct comparison with the JB55 capture data, WGS of one JB55 library was performed on a MiSeq FGx using RUO mode and paired-end (75×75 cycles) sequencing on a v3 cartridge (150 cycles).

#### 2.3.6 Sequence Data Analysis

FASTQ files were imported into the CLC Genomics Workbench v12.0.1 (QIAGEN) and all sample data were analyzed with a custom workflow. The workflow, which was applied to all samples, included five steps. First, reads were trimmed based on quality (including ambiguous bases) as well as adapter read-through (for paired reads only). Trimmed reads were then mapped to the GRCh38 human reference genome using stringent mapping parameters designed to prevent off-target mapping of exogenous DNA (e.g., bacterial DNA). Modifications to the default mapping parameters were a length fraction of 0.85 and stringency fraction of 0.95, and non-specific reads (i.e. reads that produce an equal alignment score for multiple mapping regions) were ignored. Next, duplicate mapped reads were removed with 20% maximum representation of minority sequence. Local realignment of unaligned ends was performed to improve indel alignment in the mapping. The alleles observed at each of the 5,422 FORCE SNPs was determined using the Identify Known Mutations from Mappings tool, ignoring broken pairs in the bone set sequence data. Various analysis metrics such as the total number of reads, percent mapped, and percent duplicate reads were reported throughout the workflow using different CLC Genomics Workbench quality control (e.g., QC for Targeted Sequencing) tools.

A custom Microsoft Excel (Redmond, WA) template was developed for genotype generation based on the CLC Identify Known Mutations from Mappings output. The targeted SNP had to be covered by a minimum of 10 reads for analysis. Furthermore, in order for a FORCE SNP to be called (i.e. used for downstream analyses), genotypes required allele frequency of 90% or greater to be called as a homozygote and heterozygous genotypes needed at least 30% MAF. SNPs meeting the minimum coverage requirement (10X) but with imbalanced alleles (10-30% MAF) that could not be confidently called homozygote or heterozygote were noted as “imbalanced” and excluded from further analyses. The called SNPs were separated by type (e.g., kinship SNPs, aiSNPs, piSNPs etc.) for analysis and interpretation as described below.

#### 2.3.7 SNP Concordance Assessment

The genotypes derived from the FORCE capture data of the four control DNAs (2800M, K562, NA12877 and NA12878) and one of the historical bone samples (JB55) were compared to previously published data. For 2800M, data from the ForenSeq DNA Library Preparation Kit were used for comparison [27] taking into account reverse strand orientation at 52 markers. For the remaining control DNA samples, published variant call format (VCF) files from WGS data were used for the concordance assessment ([28] for K562, [29,30] for NA12877 and NA12878). The VCF files for K562 and NA12877 included only those positions in which a variant was detected; thus, the genotype was assumed to be homozygous for the reference allele at SNPs missing from the VCF for the purpose of this comparison. A preliminary assessment identified a few issues in the published control DNA datasets that impacted concordance. There was an error in the K562 VCF for rs10892689, and the genotype for this SNP was determined by visual inspection of the associated BAM [28]. The NA12878 VCF included all SNP positions regardless of the detection of a non-reference allele. Five FORCE SNPs were missing from the NA12878 VCF, and the associated BAM [29,30] was used to confirm the genotype at those positions. Additionally, strand orientation was corrected in the published genotypes at two SNP markers (rs6670984 and rs4922532). For JB55, published genotypes produced with the Precision ID Ancestry Panel (n=165) and targeted Y-SNP typing (n=4) as described in [12] were used for comparison to the FORCE profile. The genotype with the highest coverage was utilized when discordant data were observed between the two Precision ID replicates analyzed in the previously published study. Additionally, there were 16 SNPs without sufficient coverage (<10X) in either Precision ID replicate; these markers were excluded from the concordance assessment.

Supplementary to called SNPs, “imbalanced” FORCE genotypes were also compared to the published data. Imbalanced genotypes were observed at SNPs with sufficient coverage (≥10X), but where the allele frequency thresholds for homozygote/heterozygote calling were not met. When the minor allele(s) was observed with base quality < 20 and <10% forward reverse balance, indicating sequencing error, the artifact (minor) allele was removed, and the observed genotype noted for the imbalanced SNP was modified. This was only performed for the concordance assessment. No imbalanced SNPs were utilized in downstream analyses (e.g., ancestry predictions, kinship comparisons).

When discordance was observed between the FORCE and published genotype, the FORCE BAM and the BAM associated with the published data were reviewed to confirm the genotypes at the SNP marker. If the genotypes were still inconsistent after further review, the cause of the discrepancy was investigated.

#### 2.3.8 Biogeographical ancestry predictions

Biogeographical ancestry predictions, based on the aiSNPs, were performed with two different methods: principal component analysis (PCA) and naïve Bayes analysis. An R script [24] was developed, in which the R package *prcomp* [31] was used for the PCA analysis with SNP profiles from the 1000 Genomes project [16] to train the PCA model. In the same script we included the naïve Bayes [32] analysis method by calculating the likelihood Pr(DNA|Population X), for each major continental population group (EUR, EAS, AMR, SAS and AFR). Allele frequencies from the 1000 Genomes project [16] were used for these likelihood calculations. Since the allele frequencies were missing for one aiSNP (rs10954737) in the 1000 Genomes dataset, this marker was excluded from analysis and ancestry predictions were based on 240 aiSNPs. Posterior probabilities were calculated assuming a flat prior. These previously characterized aiSNPs allow for clear separation of the major population groups EUR, AFR, EAS and SAS [33].

#### 2.3.9 Phenotype predictions

The 41 piSNPs included in the FORCE panel are consistent with the markers of the HirisPlex-S System [34]. Using the Excel template described in section 2.3.6, the genotypes from sequence data were converted into the correct allele orientation and assigned “0”, “1”, “2” or “N/A” based upon a customized R script [35]. The converted genotypes were then submitted to the HirisPlex-S System webtool for phenotype prediction [36]. The Excel template organizes the converted genotypes of the piSNPs either for easy manual entry or for batch analysis using the file upload functionality. The FORCE data were uploaded as a single file and the HirisPlex-S output was downloaded. The results were compared to self-reported eye color and hair color for the seven additional non-probative reference samples (18-24; Table 4).

#### 2.3.10 Y haplogroup predictions

Y haplogroups were predicted based upon the 829 Y-SNPs included in the FORCE panel. Called Y-SNPs were compared to the Yleaf WGS HG38 Position file [37,38] within the Excel template and variants (non-reference alleles) were denoted with the corresponding Y haplogroup. Identified haplogroups were reviewed manually and the most refined Y haplogroup was reported. The results were further inspected to ensure that the variants observed were expected based upon the predicted Y haplogroup (e.g., variants associated with ancestral haplogroups of the reported haplogroup). Though not utilized in this study, Yleaf offers automated Y haplogroup predictions [37]. Yleaf can analyze both FASTQ and BAM files; however, current functionality does not allow for duplicate removal and therefore it is recommended to utilize FORCE BAM files after mapped duplicate removal for haplogroup predictions performed with the Yleaf software.

#### 2.3.11 Kinship statistics (kinship SNP and X-SNP)

##### Pairwise kinship calculations (kinship SNPs)

Likelihoods were calculated based on observed kinship SNP data given seven different relationship hypotheses (parent/offspring, full siblings, half siblings, first cousins, first cousins once removed, second cousins and second cousins once removed) and one hypothesis representing no genetic relationship between the two compared individuals (i.e. unrelated). From these likelihoods, likelihood ratios (LRs) were calculated for each related hypothesis using the unrelated hypothesis as the alternative hypothesis (i.e. LR=Pr(DNA|H1)/Pr(DNA|H2), where H1 was the hypothesis representing one of the related relationships given above and H2 was the unrelated hypothesis). These kinship calculations were performed in a pairwise fashion between all reference samples as well as between each “Unknown” (i.e. WWII bone sample) and each reference sample. Posterior probabilities were calculated assuming a flat prior. The likelihoods were calculated using Merlin with the preferences described above (section 2.2). R was again used to format and manage input files (e.g., filter SNPs when having partial SNP profiles) as well as to handle the outputs from Merlin and for the plotting of distribution curves.

##### Pedigree kinship calculations (kinship SNPs)

In addition to the pairwise kinship calculation, kinship statistics were also calculated in a pedigree fashion. Here, SNP profiles for all reference individuals, in each reference family, were included in the likelihood calculations. In this approach each Unknown sample was tested as the missing person in each reference pedigree (Figure S1). For each such pedigree, the LR was calculated as LR=Pr(DNA|”H1: The tested Unknown sample comes from the missing person”)/Pr(DNA|”H2: The tested Unknown sample comes from an individual unrelated to the reference pedigree”). The likelihoods were calculated using Merlin with the preferences described above (section 2.2). As previously noted, a front-end script was written in R to format input files and to handle the outputs from Merlin. Kinship predictions with strong statistical support were defined as those that met the following two criteria: LR ≥ 10,000 (log 10 LR ≥ 4) *and* posterior probability ≥ 99.99% given equal priors for the tested hypotheses.

##### Kinship calculations using X-SNPs

In order to demonstrate the possibility to calculate kinship statistics solely based on X-SNPs, two samples were selected from family D: sample 12 (“Reference”) and sample 14.1 (“Unknown”). These two individuals are full siblings (Figure 2) and their X-SNP genotypes were used to calculate the LR=Pr(DNA| “Sample 12 and sample 14.1 come from two full siblings)/ Pr(DNA| “Sample 12 and sample 14.1 come from two unrelated individuals”). The X-specific version of Merlin (“MINX”) calculated the likelihoods of the two hypotheses, using European allele frequencies and genetic positions obtained from Rutgers map [16,23]. As previously noted, R was used to format input files and to handle the outputs from Merlin.

**Figure 2.**
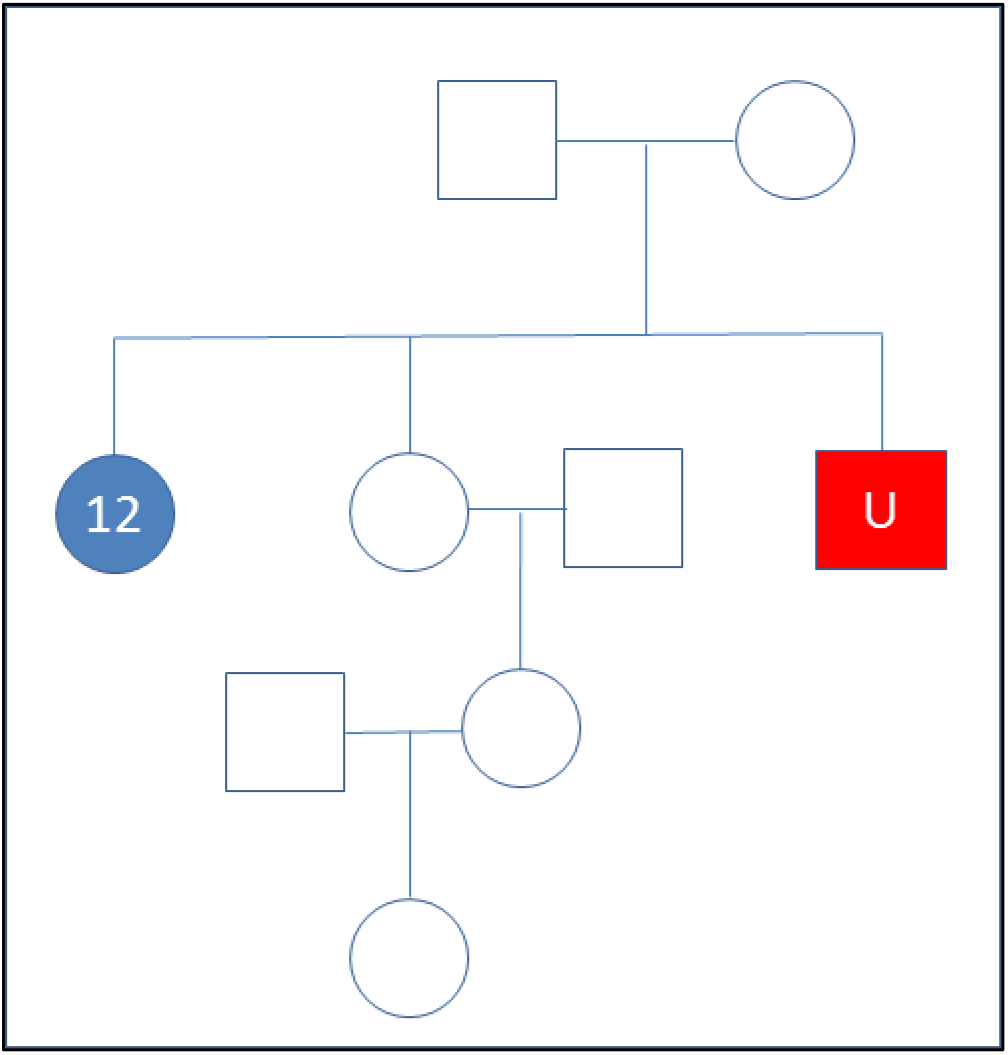
Pedigree of Unknown (sample 14.1) and the reference sample (12) from family D utilized in X-SNP kinship assessment.

## 3. Results and Discussion

### 3.1 Simulated Panel Performance

The results from the kinship simulations clearly demonstrate that the kinship SNPs could be used to solve the majority of kinship case scenarios down to first cousins, including cases with partial SNP profiles (Figure 3). Furthermore, this set of kinship SNPs can be informative in cases involving a more distant relationship issue (such as second cousins) (Figure S2); however, the LR distribution curves for such cases for true related and true unrelated cases are overlapping.

**Figure 3.**
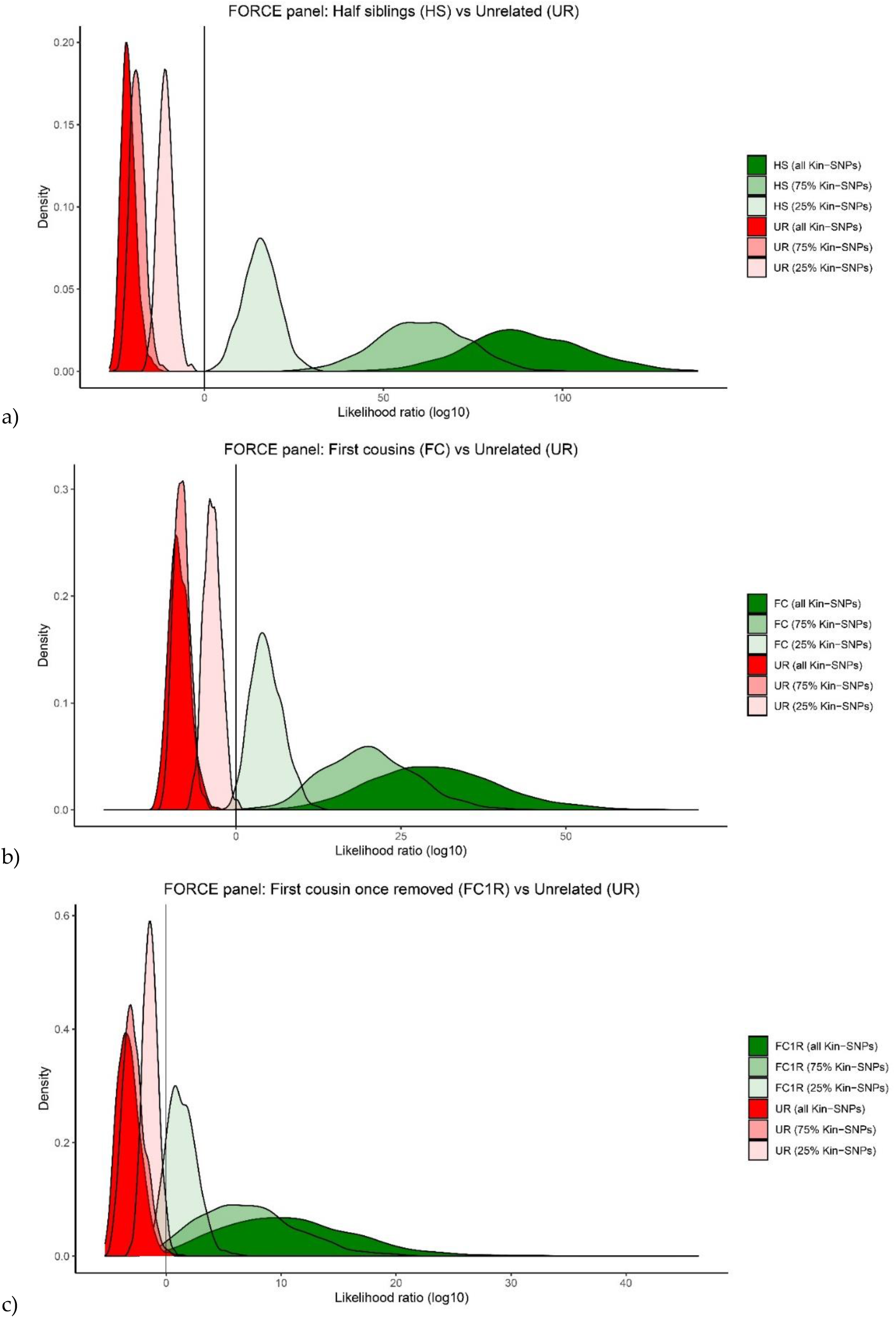
Simulation results for pairwise kinship predictions. Simulations were performed between an unknown sample at varying levels of kinship SNP recovery (100%, 75%, and 25%) compared to a complete reference sample profile. a) 2^nd^ degree (half siblings) vs. unrelated (UR); b) 3^rd^ degree (first cousins) vs. UR; c) 4^th^ degree (first cousins once removed) vs. UR. Simulation results for additional relationships (1^st^ degree and 5^th^ degree) are presented in Figure S2.

### 3.2 SNP Capture Assay Laboratory Testing

Detailed sequencing and SNP coverage metrics for all samples can be found in Table S2. The number of called SNPs, which met the 10X coverage and allele frequency requirements for homozygotes and heterozygotes, varied by sex due to the number of Y-SNPs in the assay. The maximum number of called SNPs was 5,422 for males and 4,593 for females. All reference and control DNA SNP profiles were 94.7%-99.8% complete when accounting for sex, with an average of 5,355 SNPs for males and 4,552 SNPs for females (Table 5). The average coverage of the called SNPs in reference/control samples was 225X (Table S2). The standard deviation of called SNPs in the reference/control samples was approximately 42% of the average coverage. As noted in the methods, some SNPs had fewer probes than others. Hence, average coverage by bait count was investigated (Figure S3). For autosomal SNPs having all four baits (n=4,267), the average coverage across the reference/control samples was 241X. The average coverage dropped to 180X for autosomal SNPs with three baits (n=73) and 122X for autosomal SNPs with two baits (n=6). The single autosomal (kinship) SNP (rs4847178) with one bait (right flank) produced an average coverage of 157X, with called data for 100% of the reference/control samples. All 246 X-SNPs were targeted by two baits, averaging 166X in coverage for females and 89X for males. The 812 Y-SNPs with two baits produced an average coverage of 105X in male reference/control samples, with the reduced coverage observed at the 17 Y-SNPs with only one bait (50X). There were 13 SNPs (including two kinship SNPs, one iiSNP and ten Y-SNPs) that failed to produce sufficient coverage (10X) for any sample (Table S3). An additional 17 SNPs (six kinship SNPs, one iiSNP, five X-SNPs and five Y-SNPs) had no call for a majority (>50%) of reference/control samples. Of these 30 poor performing SNPs, four had one bait that failed during the kit design. Three of these four SNPs with one failed bait were Y-SNPs (which consequently failed in all samples due to Y-SNPs only targeted by two baits per SNP). In all three cases, the failed bait was associated with the reference allele. The fourth poorly performing SNP that also had a failed bait (rs662185) had coverage and allele imbalance issues in the reference/control samples (see section 3.3 below). Other factors such as the specific baits included (e.g., reference allele, alternate allele, right flank, or left flank) and the surrounding sequence motif (e.g., insertions and deletions impacting alignment) may have affected SNP performance. As a result of these findings, the capture kit will be modified in the future to remove poor performing SNPs. The modified / “beta” FORCE kit will also eliminate baits targeting SNPs that were later discovered to lack one or more of the FORCE panel criteria (e.g., clinical relevance or triallelic SNPs).

**Table 5.**
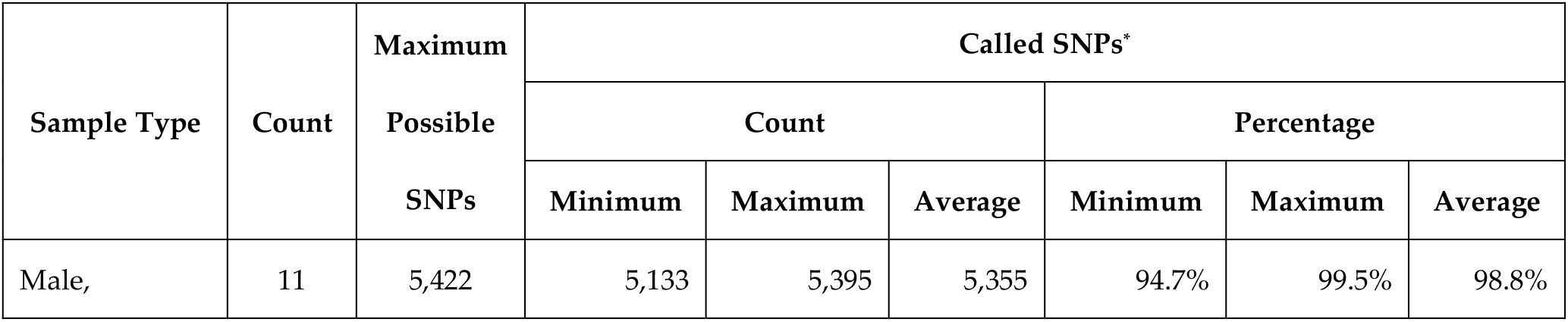
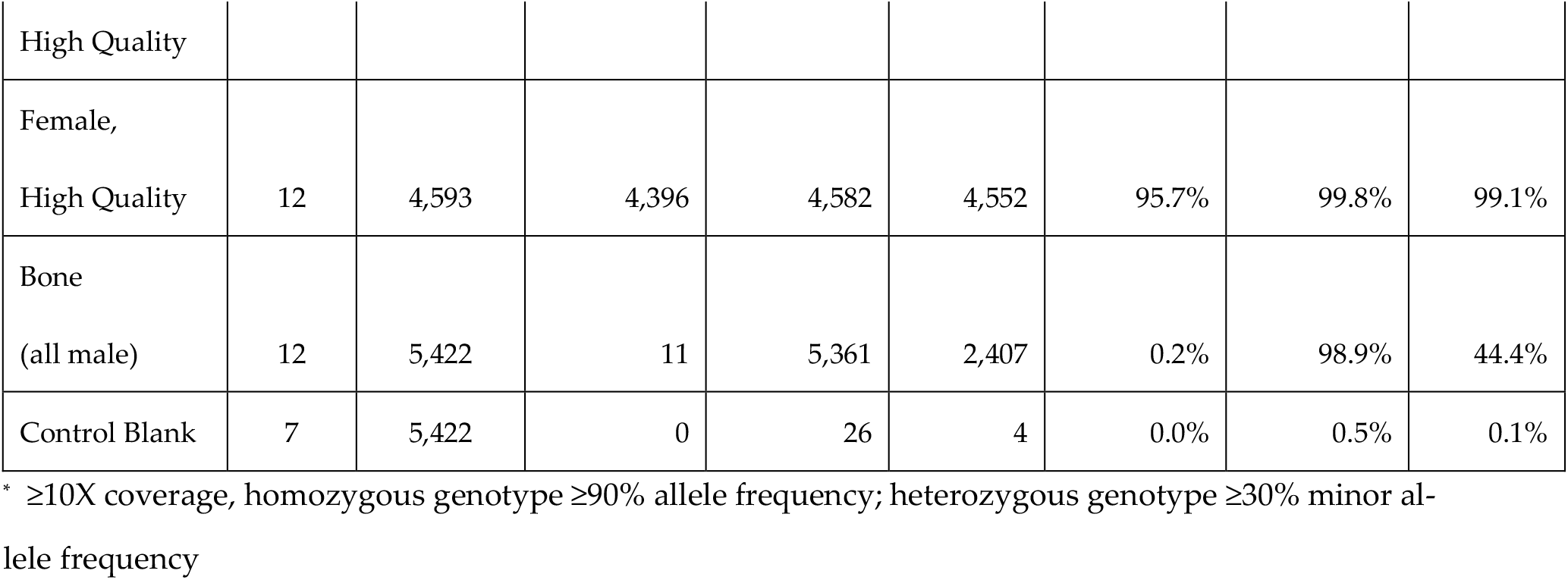
Summary of the number and proportion of called SNPs after capture of the FORCE panel targets. The results are broken down by sample type. High quality includes family references, additional non-probative reference samples, and control DNAs.

The bone sample results varied considerably, ranging from 11-5,361 called SNPs and an average of 44.4% of the possible 5,422 SNPs across the 12 DNA extracts (Table S2). To compare the number of called SNPs per bone, the results from each DNA extract were averaged for bone samples with multiple DNA extracts (Figure 5). Four of the seven bone samples produced 3,719 or more SNPs (two USS *Oklahoma* (3 and 7) and the two historical bones (JB55 and sample 26)). These four samples were extracted with either PCIA followed by MinElute purification or the Dabney DNA extraction method. In contrast, the bone samples that were extracted with QIAquick produced fewer called SNPs. The three QIAquick replicates from the Italy bone sample (14) yielded 282-5,121 called SNPs, for an average of 2,308. The Austria (11) and Tarawa (17) bone samples, which were also extracted with QIAquick, produced only 50 and 14 called SNPs on average, respectively. The Tarawa bone sample was known to be chemically treated with formalin, which was the standard postmortem procedure during WWII. Such chemically treated samples have been shown to produce poor DNA sequencing results [10]. The bone recovered from Austria may have also received the same standard WWII postmortem chemical treatment. It is important to note that the variable results obtained from the bone samples may be due to a combination of differing factors, including separate case contexts, the DNA extraction method utilized, as well as the remaining DNA extract volume available for library preparation. The sample selection strategy for this study had prioritized known, non-probative samples with available DNA extracts from both the bone sample and associated family reference samples. Therefore, it was not possible to control for DNA extract preparation and other variables that may have a considerable impact on SNP capture results. An investigation into DNA extraction methods for downstream SNP capture success is an area of future research.

**Figure 5.**
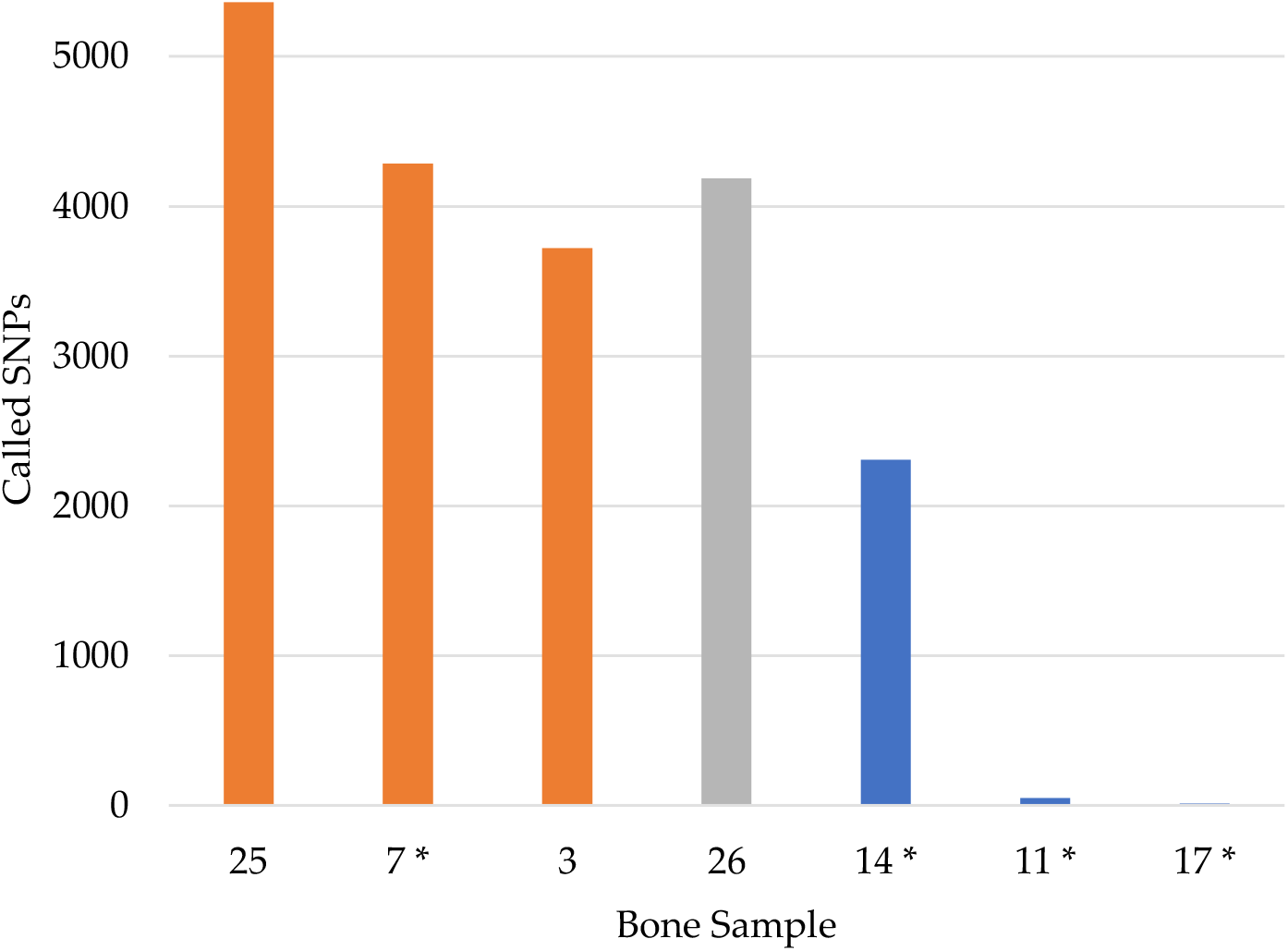
The number of called SNPs per bone sample (maximum 5,422). An asterisk (*) represents the average number of called SNPs for a bone sample with multiple DNA extracts. The color indicates the DNA extraction method: orange = phenol chloroform isoamyl alcohol (PCIA) extracts purified with MinElute; grey = Dabney; blue = QIAquick.

In the absence of capture, the bone DNA extracts produced 2-1,539 SNPs at 1X coverage and zero SNPs met the 10X calling threshold. These details and the remaining shotgun sequencing results are shown in Table S4. The proportion of reads aligning to the FORCE panel regions (i.e. mapped read specificity) was significantly increased by capture, from an average of 0.03% in the shotgun data to 11.4% in the capture data. The capture procedure thus resulted in a 380-fold increase in mapped read specificity over shotgun sequencing. Of note, shotgun sequencing of the four control DNAs produced only partial 10X FORCE SNP profiles. Thus, the capture enrichment was necessary for all samples, including references and control DNAs, in order to meet the 10X coverage requirement for SNP calling. The improved SNP profiling of the captured libraries demonstrates the efficacy of the enrichment step in obtaining on-target FORCE panel SNPs.

### 3.3 Concordance with Published SNP Data

Overall, 14,519 (99.97%) SNPs were concordant between the 14,524 called FORCE genotypes and published data of the four control DNA samples (Figure 6; Table S5). Complete concordance (100%) was observed for 2800M across the 170 SNPs overlapping called FORCE and expected ForenSeq genotypes. A high rate of concordance (99.98%) was also observed for both NA12877 and NA12878 when considering the called FORCE genotypes for each sample (5,383 and 4,576 SNPs, respectively). Though slightly lower, FORCE data of K562 produced concordant genotypes with published data at 99.93% of the 4,395 compared SNPs.

**Figure 6.**
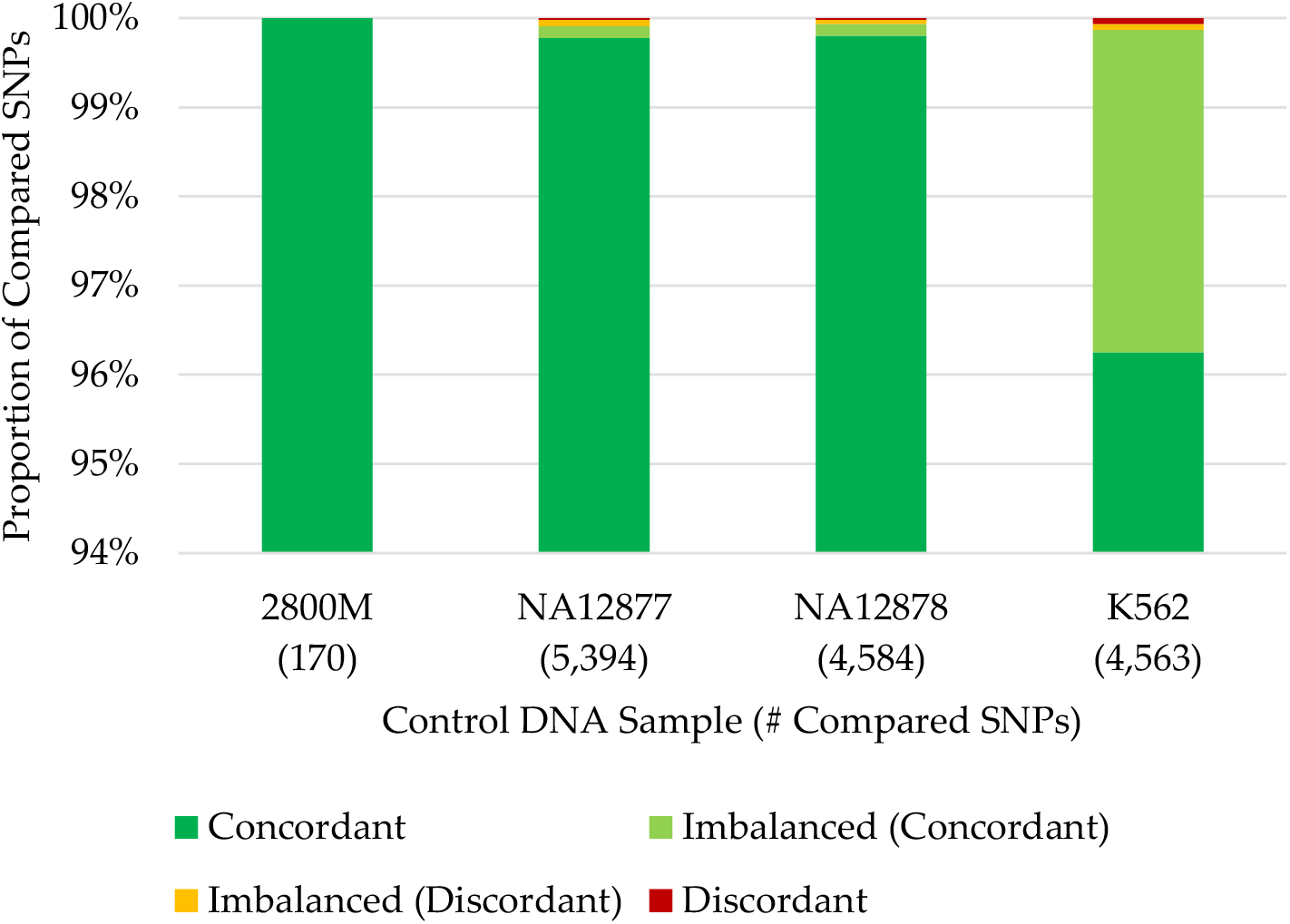
SNP concordance of FORCE capture data compared to published genotypes from the four control DNA samples.

The five discordant called FORCE genotypes were all kinship SNPs, and each discordance was observed in only one control DNA sample (Table S6). The greatest proportion of discordant calls (60%) was observed in the K562 data. Two of three discordant called SNPs produced high-coverage homozygote genotypes in the FORCE data, while heterozygote genotypes were observed in the published K562 BAM but with allele imbalance (MAF ∼20%). The third K562 discordance may be the result of incorrect genotyping based on the published data, as all reads at the SNP location in the published BAM were forward-oriented and unpaired. The greater discordance observed in the K562 data compared to the other control samples is likely due to well-known immortalization effects [28] as well as lot-to-lot variability of this cell line [39]. For NA12877, there was one difference between the called FORCE SNP and the published data at rs229352. Insufficient coverage (<10X) at this same SNP was observed for many reference/control samples (n=7). Interestingly, NA12878 and K562 were expected to be homozygous for the alternate allele (GG) at rs229352 based on the published data (this marker is not included in ForenSeq and therefore the expected genotype for 2800M is unknown). Analysis of the rs229352 bait design identified that the alternate allele (G) or left flank did not map under the stringent mapping parameters utilized in the develop workflow. Furthermore, review of the published BAM showed a 5-bp insertion located 30 bp upstream (5’) of the SNP on the reads with the G allele at rs229352. For NA12878, the discordant called SNP (rs662185) was identified as a poor performing SNP in section 3.2, with nearly half (47.8%) of the reference/control samples failing to produce sufficient coverage (<10X) and an additional 17.4% with imbalanced genotypes (Table S3). In NA12878, significantly lower coverage (10X) was observed at rs662185 compared to the average coverage of the other FORCE SNPs (>250X for called SNPs; Table S2). Similar to rs229352, the published NA12878 BAM showed a 4-bp deletion associated with the G allele at rs662185 located 5 bp downstream (3’) of the SNP. Additionally, evaluation of the three baits for this marker (the right flank bait failed during design) revealed that the alternate allele bait included in the kit was incorrect (T instead of G). Inspection of the baits for all targets identified an additional 522 markers with the incorrect alternate allele. However, these errors in the bait design appeared to have little to no impact on the SNP performance, as only seven (1.3%) of the 523 SNPs with the wrong alternate allele bait performed poorly (<75% called) in reference/control samples (Table S3). Therefore, the discordances observed at rs229352 and rs662185 are likely due to the indel clusters associated with the alternate allele, which may impact the probe hybridization efficacy and/or prevent mapping under the stringent parameters applied.

Though not utilized for other analyses in this study, the concordance of the 187 observed genotypes at imbalanced FORCE SNPs was assessed for the four control DNA samples. It is noted that 95.19% (178) of the observed imbalanced genotypes were consistent with the published data (Table S5). Unsurprisingly, a majority (168 of 187; ∼90%) of the imbalanced genotypes were detected in K562 (Table S5). As stated above, stochastic variation is expected in the K562 cell line, resulting in allele imbalance and loss of heterozygosity [28]. Furthermore, these cell line artifacts may be exacerbated by the relatively low input (1 ng) into library preparation [10]. Of the nine discordances found in the imbalanced genotypes, rs169250 accounted for three instances. The imbalanced FORCE genotypes at rs169250 were discordant for three of the control DNA samples (NA12877, NA12878 and K562; this marker is not targeted by ForenSeq and therefore no published genotype was available for 2800M). Further investigation of the BAMs revealed that rs169250 is located in the middle of a 12-bp polycytosine stretch, and thus the complexity of the region for sequencing and/or alignment likely caused discordant genotypes. As a result, future designs of the FORCE panel may exclude rs169250 to ensure successful and accurate genotyping. The remaining imbalanced FORCE genotypes for NA12877 (n=3) and NA12878 (n=1) that were discordant with the published data appear to be the result of non-specific mapping in the FORCE data. The last two discordances involving imbalanced FORCE genotypes were found in K562. One appeared to be the result of non-specific mapping in the published BAM (rs241408), while the other (rs7117433) could not be resolved and may be due to cell line artifacts previously discussed.

In the end, considering both called and imbalanced FORCE genotypes, a total of 99.90% (14,697 out of 14,711) SNPs were in agreement between the FORCE and published control DNA datasets (Table S5). There were, however, a total of 69 genotypes could not be compared out of a maximum of 14,780 possible SNPs (Table S5) because no call was possible for the FORCE or published data (<10X). Most (68) of the failed SNPs were observed in the FORCE data, while one SNP for K562 had low coverage (5X) in the published BAM that prevented the associated published genotype from being confirmed. The number of failed SNPs varied by sample: two of 172 SNPs for 2800M, nine of 4,593 SNPs for NA12878, 28 of 5,422 SNPs for NA12877 and 30 of 4,593 SNPs for K562. Of note, FORCE data from all four control DNA samples failed to produce called genotypes for 16 SNPs. Thirteen of these SNPs failed to produce sufficient coverage for any samples (Table S3), as previously noted in section 3.2. The three additional poor performing SNPs included two X-SNPs and one Y-SNP. Most (∼70%) reference samples failed to produce called genotypes at the two X-SNPs, with no called data in the bone samples. The Y-SNP produced greater reference/control samples success (>65% of male references/controls with a called genotype), but 0% SNP recovery for the bone samples.

Published and FORCE data for JB55, the only bone sample evaluated for concordance, enabled the comparison of 153 SNPs. Sixteen of the 169 overlapping SNPs were excluded from comparison due to insufficient coverage (<10X) in the published Precision ID data. JB55 had a concordance rate of 98.69% with 151 of 153 concordant SNPs, and two discordant aiSNPs were observed (Table S6). The FORCE genotype (CC) for the first discordant marker (rs1879488) was inconsistent with the highest coverage Precision ID replicate from the published study (AC with 395X in replicate 2); however, the FORCE results were concordant with the replicate with slightly lower coverage (CC with 236X in replicate 1). The second difference was observed at a SNP with low coverage in both Precision ID replicates (8X and 16X), potentially the result of non-specific amplification and/or mapping. In the case of these two discordant SNPs, stricter parameters such as a higher coverage threshold or heterozygote threshold than what was used for the previous JB55 study (≥6X and ≥10% MAF) [12] may have improved the concordance rate for JB55.

### 3.4 Ancestry, phenotype, and Y haplogroup predictions

The completeness of the reference and control sample profiles allowed for ancestry, phenotype, and Y-chromosomal haplogroup predictions, which were consistent with available metadata (Table S7). The predictive results from the bone sample DNA extracts were much more variable due to the wide range of SNPs obtained. As an example of the bone sample data, the ancestry of sample 7.2 was predicted to be 100% European from 210 called aiSNPs (Figure 7). Phenotype predictions were also possible for this bone sample (7.2) using 33 called piSNPs, which estimated blue eyes (93.2% probability), blond-brown hair (66.0% and 27.4% probabilities, respectively), light hair shade (92.8% probability), and pale-intermediate skin (54.1% and 41.6% probabilities, respectively). The Y haplogroup was predicted to be R1a1a1b1a2b∼ from 370 Y-SNPs, which was the same Y haplogroup that was predicted for the paternal nephew (sample 4). These predictions were possible even with the partial SNP profile (4,165, or 76.8%, of possible 5,422 called SNPs) generated from this degraded USS *Oklahoma* sample. Thus, hybridization capture was effective in producing sufficient SNPs from the multiple marker types included within the FORCE panel to allow for predictive information to be gained.

**Figure 7.**
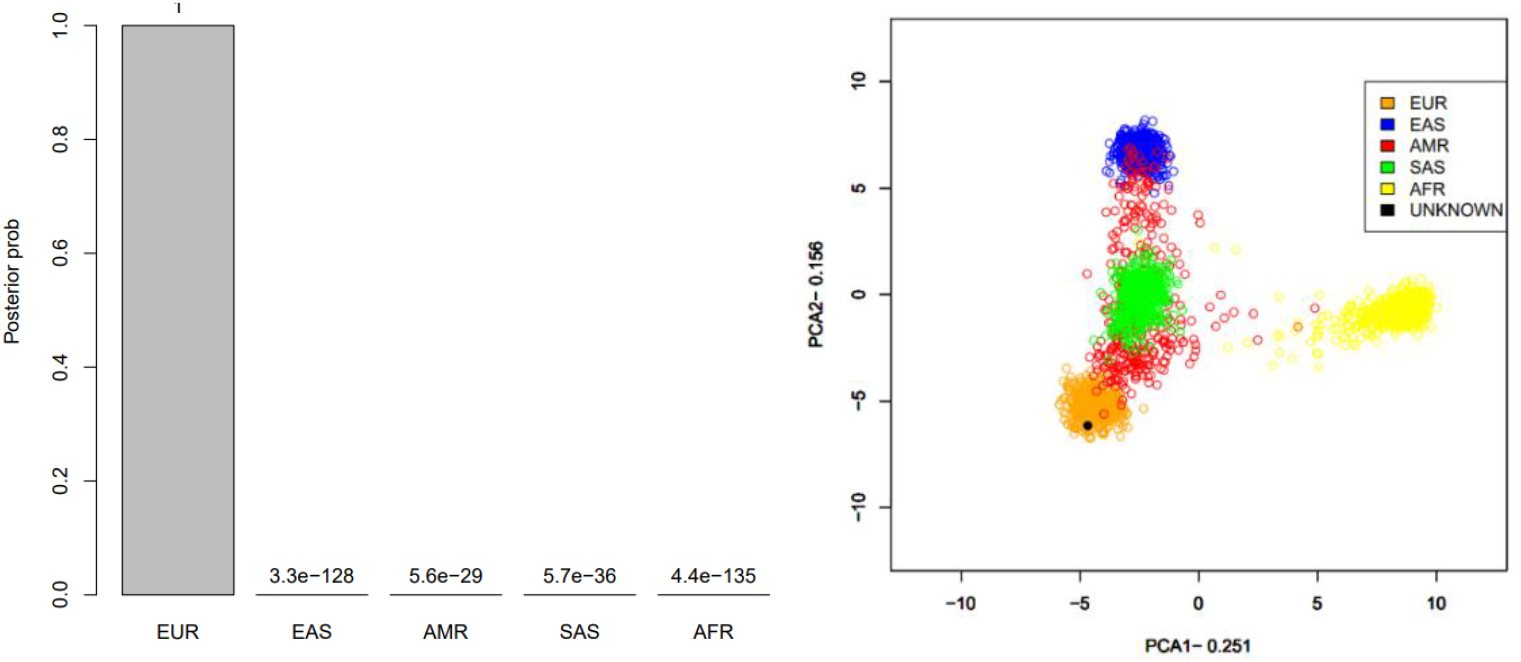
An illustration of ancestry predictions, here exemplified with predictions for sample 7.2. Individuals from the 1000 Genomes project (grouped into the five different continental groups: EUR (European), EAS (East Asian), AMR (American), SAS (South Asian) and AFR (African)) were used as reference data. Left = Posterior probabilities using a naïve Bayes approach; Right = Prediction using a PCA.

### 3.5 Kinship Predictions

The related reference to reference sample kinship prediction results are shown in Figure 8. The 1^st^ degree (parent/offspring and full sibling) predictions produced log 10 LRs from 287.7-439.3 with well over 99.99% posterior probabilities, demonstrating extremely strong statistical support of these predictions. The log 10 LRs for the 3^rd^ degree reference to reference predictions were 34.4 and 37.8 with over 99.99% posterior probability, indicating very strong statistical support. The two 5^th^ degree relationship predictions produced log 10 LRs of 3.8 and 6.5, with 99.98% and 99.99% posterior probabilities, respectively. Thus, the kinship analysis was capable of correctly predicting 5^th^ degree relatives from reference quality FORCE panel profiles. The two 6^th^ degree relationship predictions were supported with over 99% posterior probability, but with log 10 LRs of 2.2 and 2.6, indicating moderate statistical support. Considering the four examples of 5^th^-6^th^ degree relationships, log 10 LRs corresponding to moderate statistical support [40] were obtained in a majority (75%) of the 5^th^-6^th^ degree comparisons. Interestingly, sample 4 (5^th^ degree comparison) had the lowest SNP recovery of the references (95%) and sample 8 (6^th^ degree comparison) also had reduced SNP recovery (98%). All but one of the remaining references had >99% complete FORCE profiles (sample 13 at 98%). Therefore the ability to obtain stronger statistical support for these more distant degrees of relatedness (as in the case of samples 4 and 8) could be possible when complete FORCE panel profiles are obtained. Regardless, the simulation studies indicated considerable overlap in expected versus unrelated relationships when testing 5^th^ degree relatives and beyond. When pairwise kinship predictions were performed in a blind search fashion allowing up to either 3^rd^ degree or 6^th^ degree relatedness, zero incorrect (false positive) relationships with strong statistical support were observed (Table S8).

**Figure 8.**
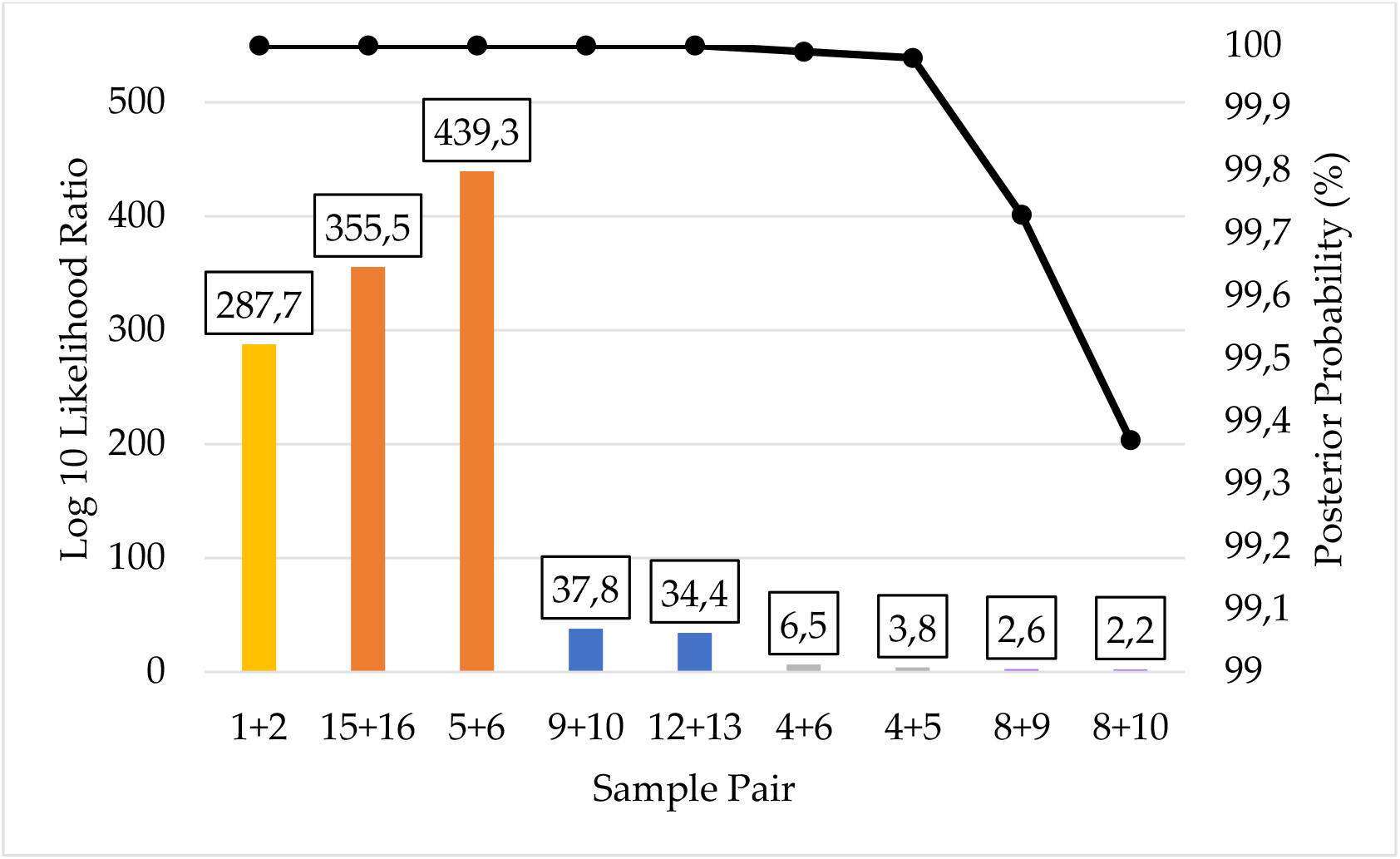
Pairwise kinship prediction log 10 likelihood ratio (LR) (left y-axis; bars) and posterior probabilities (right y-axis; black line) of reference to reference comparisons. All predictions compared the expected (known) relationships vs. unrelated. Parent/offspring = yellow (sample pair 1+2); full siblings = orange (sample pairs 15+16 and 5+6); 3^rd^ degree = blue (sample pairs 9+10 and 12+13); 5^th^ degree = grey (sample pairs 4+6 and 4+5); and 6^th^ degree = purple (sample pairs 8+9 and 8+10).

Unknown to reference pairwise kinship predictions from bone sample DNA extracts that produced 1,000 (∼25%) or more kinship SNPs are presented in Figure 9. Similar to what was observed in the reference to reference pairwise comparisons, the Unknown to reference full sibling predictions had extremely strong statistical support. The log 10 LRs of these 1^st^ degree relationship predictions were 120.7 and 492.8, with posterior probabilities over 99.99%. The three 2^nd^ degree relationship predictions produced log 10 LRs between 65.4 and 82.3 and posterior probabilities over 99.99%, also indicating extremely strong statistical support. Two of the three 3^rd^ degree relationship predictions (for sample pairs 3+1 and 14.1+13) were consistent with the reference to reference predictions. These two had log 10 LRs of 32.8 and 41.1 with posterior probabilities over 99.99%, indicating extremely strong statistical support. The other 3^rd^ degree prediction (14.2+13) also had a posterior probability over 99.99% but with a log 10 LR of 5.5, which was much lower than that of the other two 3^rd^ degree predictions. This LR indicating strong statistical support was made from a bone sample profile with only 1,324 (33.7%) kinship SNPs. It is likely that a higher LR would have been obtained if sample 14.2 had greater SNP recovery. The four 4^th^ degree relationship predictions had log 10 LRs from 4.5 to 11.8, and all had posterior probabilities over 99.99%. These 4^th^ degree Unknown to reference predictions would be classified as having strong to very strong support.

**Figure 9.**
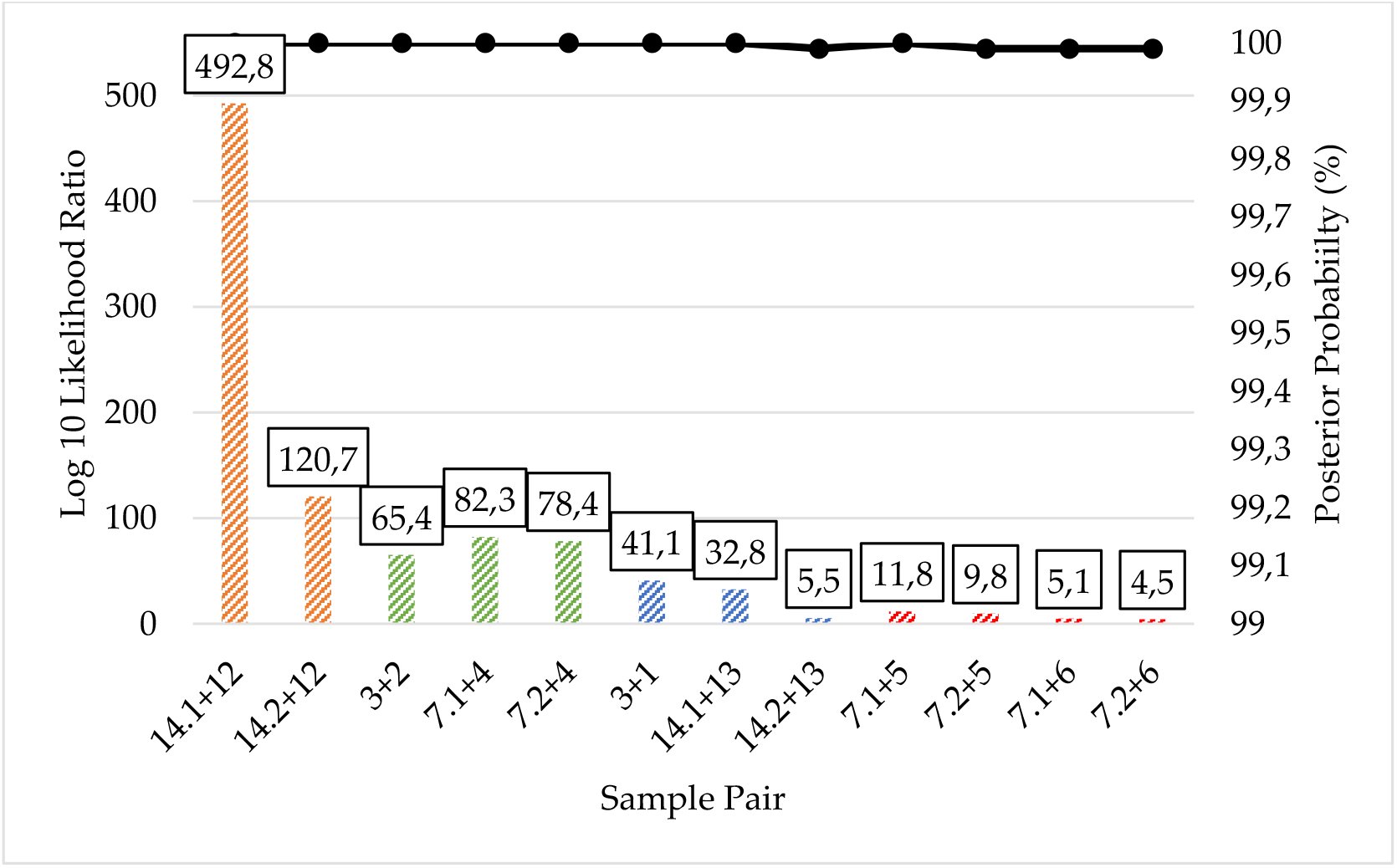
Pairwise kinship prediction log 10 likelihood ratio (LR) (left y-axis; bars) and posterior probabilities (right y-axis; black line) of Unknown to reference comparisons. Results are presented for each bone sample DNA extract that produced 1,000 (∼25%) or more kinship SNPs. All predictions compared the expected (known) relationships vs. unrelated. Full siblings = orange (sample pairs 14.1+12 and 14.2+12); 2^nd^ degree = green (sample pairs 3+2, 7.1+4, and 7.2+4); 3^rd^ degree = blue (sample pairs 3+1, 14.1+13, and 14.2+13); 4^th^ degree = red (sample pairs 7.1+5, 7.2+5, 7.1+6, and 7.2+6).

The five bone sample DNA profiles with >25% kinship SNPs were utilized for kinship predictions in a pedigree scenario, as presented in Figure 10. All of the predicted relationships indicated relatedness between the Unknown and the expected family members. These pedigree scenario relationship predictions were supported by log 10 LRs greater than 91.3 (and up to 503.7) with posterior probabilities exceeding 99.99%. Thus, the pedigree scenarios yielded extremely strong statistical support when bone sample profiles yielded ∼25% or more kinship SNPs.

**Figure 10.**
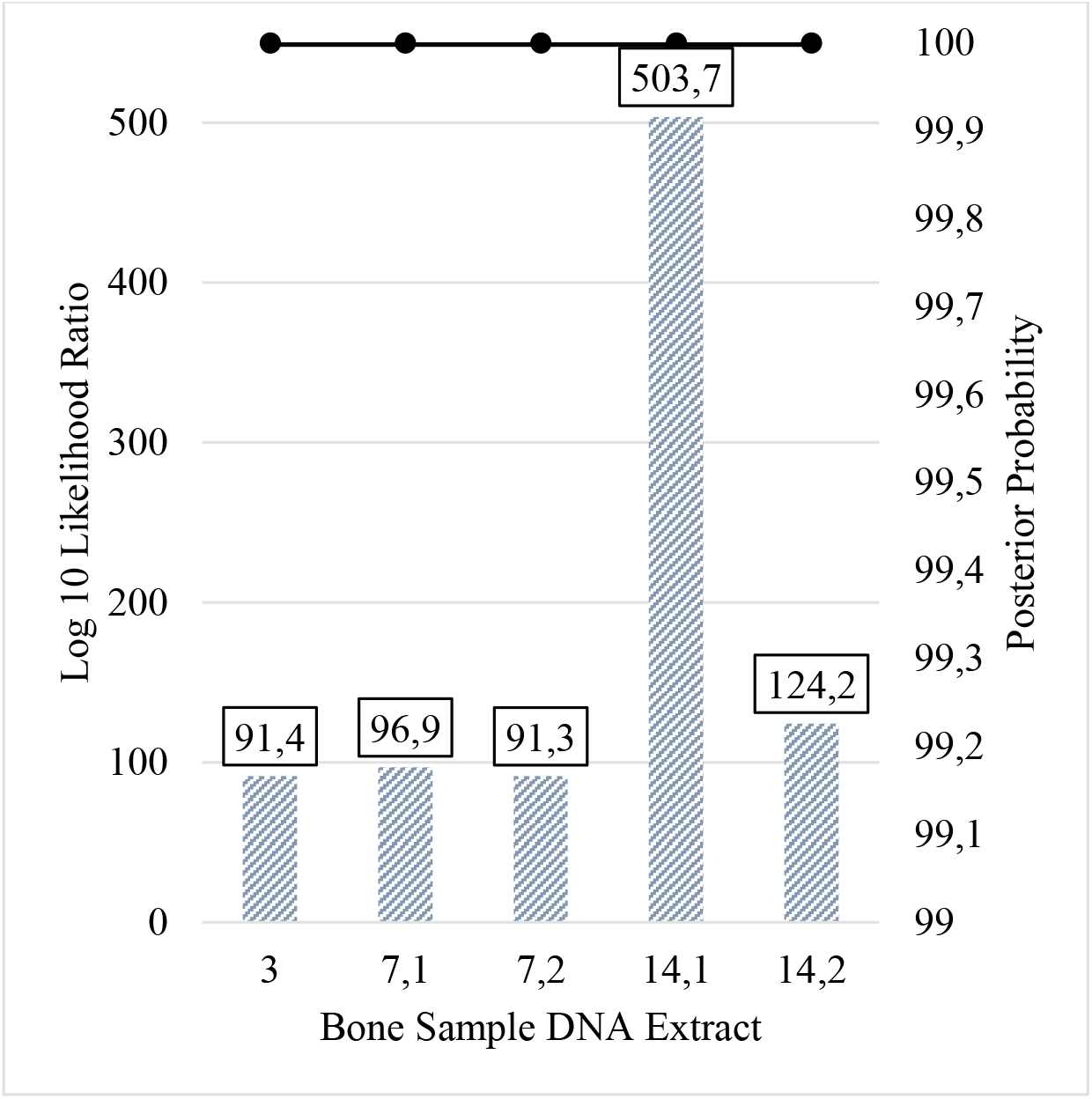
Pedigree scenario kinship prediction results of all bone samples with more than 1,000 (∼25%) kinship SNPs. All predictions compared the Unknown when situated in the expected (known) pedigree vs. unrelated. The log 10 likelihood ratio (LR) (left y-axis; bars) and posterior probabilities (right y-axis; black line) are shown.

When bone sample profiles produced minimal SNP data (10-247 kinship SNPs), the kinship predictions were expectedly weaker overall (Figure 11). Only two bone sample profiles yielded predictions that would indicate strong statistical support, and both involved 1^st^ degree relationships. Unknown (bone) sample 11.2 had 73 kinship SNPs, and its parent/offspring relationship with sample 10 was strongly supported by a log 10 LR of 5.2 and a posterior probability > 99.99%. Sample 14.3 had 247 kinship SNPs, and its full sibling relationship with sample 12 was very strongly supported by a log 10 LR of 15.8 and posterior probability > 99.99%. The remaining six relationship pairs shown in Figure 11 had low log 10 LRs (1.2 or less) and posterior probabilities (< 95%). The pedigree scenario kinship predictions did not significantly improve the log 10 LRs or posterior probabilities of the bone sample profiles with 10-247 called SNPs (Figure 12). Additionally, samples 17.1 and 17.2 yielded <15 kinship SNPs each, with some SNPs that were inconsistent with the profiles of the associated child reference (one son and one daughter of the Unknown) in these parent/offspring relationships. As a result of the poor SNP recovery, bone samples 17.1 and 17.2 were excluded altogether from the pairwise and pedigree scenario kinship predictions. Regardless of the number of kinship SNPs in the bone sample profiles, there were no incorrect (false positive) relationship predictions between Unknowns and unrelated reference samples with strong statistical support in blind search scenarios (Table S9).

**Figure 11.**
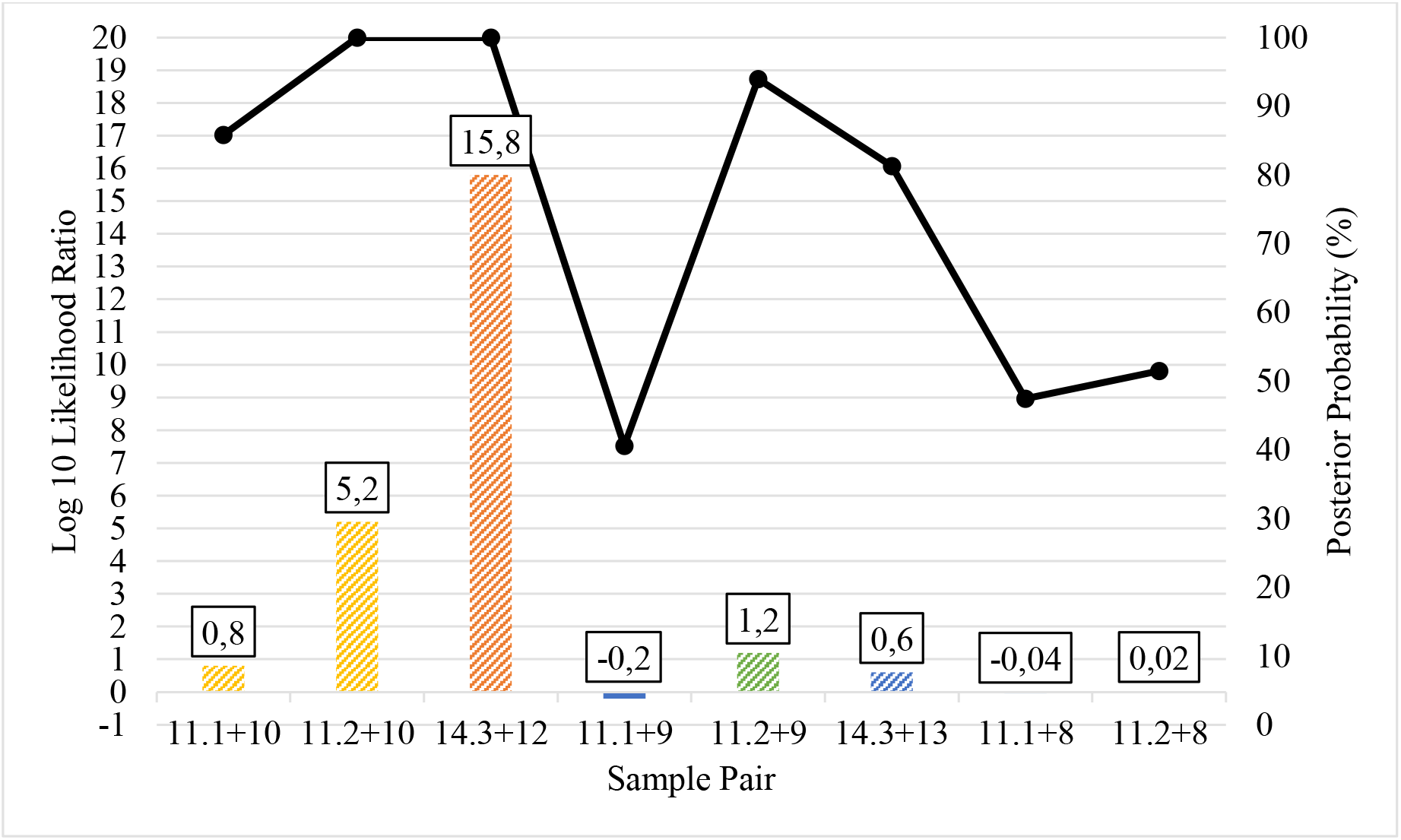
Pairwise kinship prediction log 10 likelihood ratio (LR) (left y-axis; bars) and posterior probabilities (right y-axis; black line) of Unknown to reference comparisons. Results are presented for each bone sample DNA extract that produced 10-247 kinship SNPs (all results <25% called SNPs). Samples 17.1 and 17.2 yielded <15 kinship SNP calls that were inconsistent with the expected parent/offspring relationships; therefore 17.1 and 17.2 and were excluded altogether. All predictions compared the expected (known) relationships vs. unrelated. Parent/offspring = yellow (sample pairs 11.1+10 and 11.2+10), full siblings = orange (sample pair 14.3+12); 2^nd^ degree = green (sample pairs 11.1+9 and 11.2+9); 3^rd^ degree = blue (sample pair 14.3+13); and 5^th^ degree = grey (sample pairs 11.1+8 and 11.2+8).

**Figure 12.**
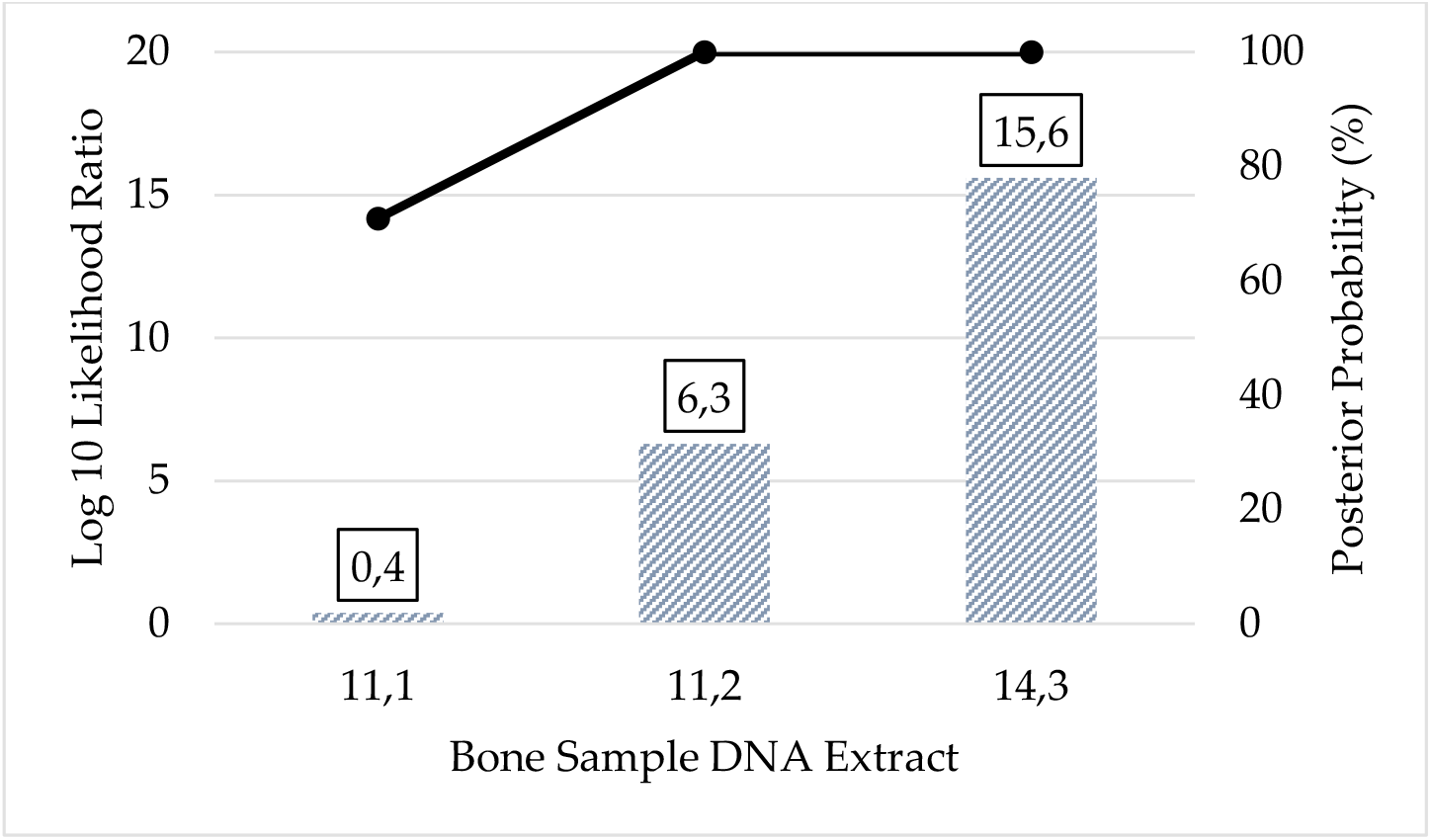
Pedigree scenario kinship prediction results of bone sample DNA extracts that produced 10-247 kinship SNPs. All predictions compared the expected (known) pedigree vs. unrelated. The log 10 likelihood ratio (LR) (left y-axis; bars) and posterior probabilities (right y-axis; black line) are shown. Samples 17.1 and 17.2 yielded <15 kinship SNP calls that were inconsistent with the expected parent/offspring relationships; therefore 17.1 and 17.2 and were excluded altogether.

Supplementary to the autosomal SNPs, 246 X-SNPs were targeted in the panel for kinship testing in specific case contexts. The X-SNPs included in the panel were used to connect individual 12 with the remains from individual 14.1 in family D. The log 10 LR (for them being full siblings versus being unrelated) was calculated to be 6.7 and a posterior probability > 99.99%; thus the X-SNP data showed to be clearly informative in the case example. The use of X-chromosomal data may be especially informative in kinship testing due to its particular mode of inheritance. For example, when comparing two alleged paternal half-sisters or an alleged paternal grandmother and a granddaughter, the exclusion power for X-chromosomal markers are not zero, in contrast to autosomal markers [41].

## 5. Conclusions

This study demonstrates the utility of the 5,422-SNP FORCE panel for forensic applications. This panel was designed to exclude clinically relevant markers to alleviate privacy concerns with DNA databanking. The SNPs included in the FORCE panel are found on microarray genotyping chips used for genetic genealogy, enabling cross-compatibility with externally produced data that will be useful in validation studies. Moreover, large reference databases are available for all of the markers included in the kit, unlike other marker types such as microhaplotypes. The FORCE panel includes kinship SNPs with forensic iiSNPs, aiSNPs, piSNPs, Y-SNPs, and X-SNPs. The hybridization capture method was capable of recovering SNP profiles from high quality reference and historical bone samples alike. A high degree of concordance (>99.9%) was observed when comparing FORCE profiles with previously published data. The few underperforming SNPs that were identified in the present study can be removed from future designs of the FORCE panel to improve efficiency. Ancestry and phenotype predictions were possible and consistent with self-reported data from reference sample donors. Additionally, high-resolution Y-haplogroup prediction was possible from 829 Y-SNPs. The novel kinship marker set of 3,931 SNPs was shown to be effective for 4^th^ degree relationship predictions with strong statistical support. Furthermore, known case examples involving previously identified WWII service members and associated family reference samples resulted in kinship predictions consistent with expected relationships. Future efforts involving the FORCE panel will entail an inter-laboratory study incorporating alternate enrichment methods for SNP profiling. This will be pertinent before the FORCE panel can be used in routine casework.

## Disclaimer

The assertions herein are those of the authors and do not necessarily represent the official position of the United States Department of Defense, the Defense Health Agency, or its entities including the Armed Forces Medical Examiner System. Any mention of commercial products was done for scientific transparency and should not be viewed as an endorsement of the product or manufacturer.

## Supplementary Materials

The following are available online at www.mdpi.com/xxx/s1, Table S1: The 5,446 SNPs targeted by the alpha version of the custom FORCE capture panel;; Table S2: Detailed sequencing and SNP coverage metrics after FORCE capture; Table S3: SNPs with called FORCE genotypes for less than 75% of the reference/control samples; Table S4: Shotgun sequencing and SNP coverage metrics; Table S5: Concordance of FORCE genotypes and published genotypes for the four control DNA samples as well as historical bone sample JB55; Table S6: Discordances observed between the FORCE and published genotypes of the four control DNA samples and historical bone sample JB55; Table S7: Ancestry, phenotype, and Y-chromosomal haplogroup predictions of reference and control DNA samples; Table S8: Blind search reference to reference pairwise comparison results; Table S9: Blind search Unknown to reference pairwise comparison results; Figure S1: Pedigree scenarios evaluated for the five WWII cases; Figure S2: Additional kinship simulations; Figure S3: SNP coverage based on the number of baits.

## Author Contributions

Conceptualization, A.T., K.S.A., and C.M.; kinship SNP selection and simulations, A.T.; hybridization capture panel design, K.S.A. in collaboration with Arbor Biosciences; hybridization capture data generation, J.D.H. and J.T.T.; formal analysis, A.T., K.S.A., and C.M.; writing—original draft preparation, A.T., K.S.A., J.D.H., and C.M.; writing—review and editing, all authors. All authors have read and agreed to the published version of the manuscript.

## Institutional Review Board Statement

The study was conducted according to the guidelines of the Declaration of Helsinki, and approved by the Defense Health Agency Office of Research Protections on November 20, 2020 (Protocol # DHQ-20-2073).

## Informed Consent Statement

All living sample donors provided informed consent for their samples to be used in research and quality improvement activities.

## Data Availability Statement

Data are stored at the AFMES-AFDIL and may be made available to approved laboratories upon written request.

## Acknowledgments

The authors would like to thank Dr. Timothy McMahon (AFMES-AFDIL) for administrative support and resources; Ursula Zipperer for anonymizing samples; Cassandra R. Taylor for assistance with laboratory experiments.

